# Vessel-on-Chip model of the microcirculation in Abdominal aortic aneurysms

**DOI:** 10.1101/2025.09.10.675261

**Authors:** Philipp C. Hauger, Karlijn B. Rombouts, Marc Vila Cuenca, Albert van Wijk, Max C. Overboom, Jan Willem Buikema, Kak Khee Yeung, Valeria V. Orlova, Peter L. Hordijk

**Affiliations:** Department of Physiology, Amsterdam University Medical Centers, Amsterdam, The Netherlands; Department of Anatomy and Embryology, Leiden University Medical Center; Department of Surgery, Amsterdam University Medical Centers, Amsterdam, location University of Amsterdam, The Netherlands; Department of Cardiology, Amsterdam University Medical Centers, Amsterdam, The Netherlands; Amsterdam Cardiovascular Sciences, Atherosclerosis and Aortic diseases, Amsterdam, The Netherlands

**Author notes:** Corresponding author: Name: Prof Peter L. Hordijk Address: Department of Physiology, O2 gebouw, De Boelelaan 1108, 1081 HZ Amsterdam.

## Abstract

Abdominal aortic aneurysms (AAA) are pathological dilations of the abdominal aorta. To date, surgical intervention is the only option for managing large AAAs, with no pharmacological therapies to prevent growth of small aneurysms. A current limitation in investigating further pharmacological avenues is the translatability of results from animal models, or from patient trials that are limited by co-morbidities and disease severity. To bridge this knowledge gap, we created a novel, patient-specific vessel-on-chip (VoC) model of the microcirculation in AAA (AAA-VoC). We found that co-culture of both C (control)-VSMCs and AAA-patient derived VSMCs with healthy, hiPSC-derived ECs generate lumenized and perfusable microvascular networks. We show that AAA-VoCs are characterized by an enlarged average vascular diameter. We furthermore found that AAA-VSMCs show phenotypical deviations from C- VSMCs after 7 days in co-culture such as increased number and surface area, indicative of a preserved pathological phenotype in our *in vitro* model. Lastly, we demonstrate that AAA-VoCs showed an increased level of pro-inflammatory cytokine expression over C-VoCs and displayed an impaired endothelial barrier function, resulting in vascular leakage. With this study, we show that AAA-VSMCs affect microvascular networks formed by healthy hiPSC-ECs and that a AAA phenotype is preserved in 3D co-culture, making this model valuable for future studies investigating treatments for AAA.

## Introduction

Abdominal aortic aneurysms (AAA) are pathological dilations of the aorta in the abdomen. AAA typically remain asymptomatic until rupture, which is associated with an overall mortality rate of <80% (1). AAA patients with a known risk of rupture, currently predicted based on aortic diameter, are treated with surgical procedures. Besides surgical intervention of large aneurysms, there is currently no therapeutical option to treat small or asymptomatic aneurysms (2). Pharmacological interventions are currently under investigation, but to date have been unsuccessful in slowing, stopping, or reducing aneurysm progression (3–5).

The lack of current pharmacological advances has raised the question for translatability of results obtained in animal models and in human observational trials and genetic studies (5). A significant limitation of animal models for AAA is that unlike humans, animals do not spontaneously develop AAA due to lifestyle factors alone, necessitating artificial disease induction methods that accelerate disease onset and introduce confounding effects related to the intervention itself (6). Furthermore, human clinical trials are limited by low and medically fragile patient populations, high rate of co-morbidities, AAA growth progression and variability, and have so far failed to produce meaningful pharmaceutical outcomes (2, 7–10).

*In vitro* cell culture systems offer strong potential to generate translatable results by utilizing human patient-derived material in a controlled environment. The aortic wall predominantly consists of endothelial cells (ECs) and vascular smooth muscle cells (VSMCs), and VSMC dysregulation plays a critical role in AAA initiation and progression (11). Moreover, the aortic wall is perfused by a microvascular network (vasa vasorum) and growing evidence suggests a role of the latter in AAA pathology (12–15), highlighting EC and VSMC dysregulation as central to disease development.

Multi-cellular approaches have been developed to study cellular crosstalk in AAA, by seeding cells on bioengineered scaffolds or ECM-like gels (16, 17). However, these techniques rely on defined structural matrices and tissue engineering geometries, constraining cells from self-organizing. This may limit their ability to recapitulate key morphogenetic processes and microenvironmental cues.

Increasing complexity in *in vitro* tissue engineering enabled the use of self-organized 3D Vessel-on-a-Chip (VoC) models, in which ECs and VSMCs are embedded in a hydrogel as single cell suspension after which they form, over the course of several days, a vascular network with perfusable lumen (18). Establishing a well-defined, self-organizing VoC model that mimics the microcirculation of the AAA vessel wall could significantly enhance our understanding of the role of ECs and VSMCs in the context of AAA pathophysiology, but such an AAA disease VoC model is currently unavailable.

In the present study, we created a novel, patient-specific AAA-VoC model mimicking a vasa vasorum-derived microcirculation, by co-culturing primary VSMCs derived from AAA patients (AAA-VSMC) or healthy individuals (C-VSMC) with healthy human induced pluripotent stem cell-derived EC (hiPSC-ECs) (Fig. 1A) to investigate the effect of patient derived VSMCs on microvascular networks of a healthy background. We show that AAA-VSMCs affect microvascular morphology and hiPSC-EC function, and that several aspects of the AAA phenotype are preserved in 3D co-culture, making this novel AAA-VoC model valuable for future studies investigating alternative treatment avenues for AAA.

**Figure 1.**
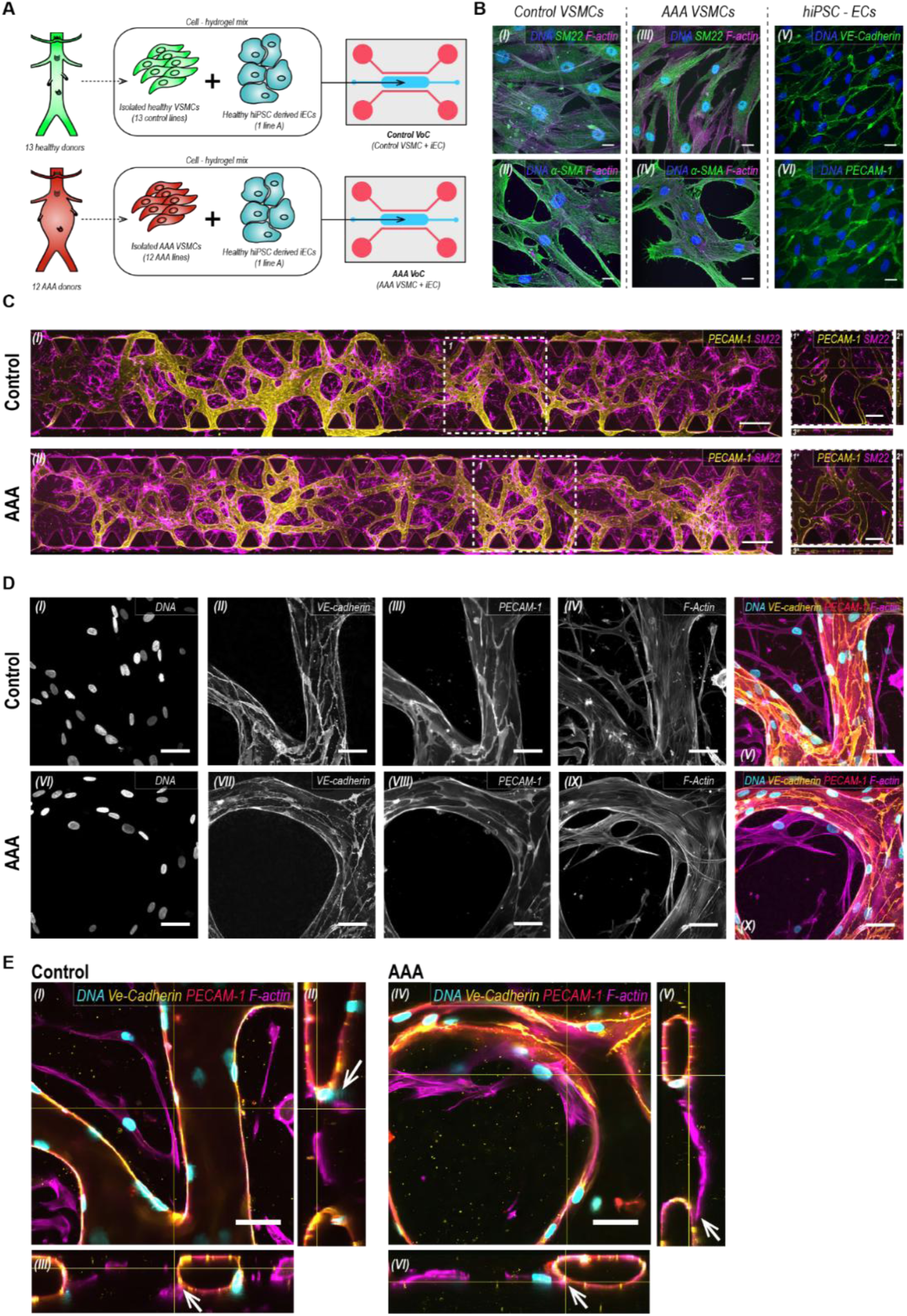
Characterization of the self-organizing VoC models. (A) Schematic overview of VSMC isolation and experimental set-up. Primary VSMCs, hiPSC-ECs and hydrogel are located in the middle channel of the microfluidic unit indicated in blue, top and bottom flanking channels contain cell culture media, indicated in red. (B) Representative confocal images of primary VSMCs immunostained for SM22 and aSMA in control (I+II) and AAA VSMCs (III+IV) (blue: DAPI, green: SM22 (I+III) / aSMA (I+IV), magenta: F-actin), and hiPSC-ECs immunostained for the endothelial markers VE-cadherin and PECAM-1 (V+VI) (blue: DAPI, green: VE-Cadherin (V) / PECAM-1 (IV), magenta: F-actin). Scale bars: 20 µm. (C) Representative confocal images of microvascular networks showing hiPSC-ECs (yellow: PECAM-1) and primary VSMCs (magenta: SM22) across the complete length of the microfluidic channel. Upper panel C-VoC, lower panel AAA-VoC. Scale bars: 500 μm. (1*) Cross- sectional images in xyz, (2*) xy and (3*) yz from a representative area (1). Scale bars: 250 µm. (D) Representative confocal images of microvascular structures in detail, upper panel in C-VoC, lower panel in AAA-VoC. (I + VI) Cell nuclei, (II + VII) VE- Cadherin, (III + VIII) PECAM-1 and (IV + IX) F-actin. (V) Composite images for C-VoC and (X) AAA-VoC (blue: DAPI, yellow: VE-Cadherin, red: PECAM-1, magenta: F-actin). Scale bars: 50 µm. (E) Representative confocal orthogonal images of (I) C-VoC in xyz and (IV) AAA-VoC in xyz depicting contact points between primary VSMCs and hiPSCs in VoC culture (contact points indicated by white arrows in xy and yz slices, C-VoC: II + III, AAA-VoC: V + VI), (blue: DAPI, yellow: VE-Cadherin, red: PECAM-1, magenta: F-actin). Scale bars: 50 µm.

## Methods

### Patient population, aortic tissue collection and primary VSMC isolation

Biopsies at the largest diameter of the aorta were obtained from 12 AAA patients during open aneurysm repair in the Amsterdam University Medical Center, location VUmc and AMC, the Netherlands. All patients were over 18 years of age and signed informed consent to participate in the study (patient characteristics: Tab. T1). Control aortic biopsies were obtained from non-dilated infrarenal aorta of 13 post-mortem kidney donors. Biopsies were transported directly after collection on ice-cold sterile 0.9% NaCl solution from the operating room to the laboratory. Tissue collection was in accordance with the regulations of the WMA Declaration of Helsinki and institutional guidelines of the Medical Ethical Committee of the VU Medical Center (Biobank 2017.121: Aortic Aneurysms, Atherosclerosis and Biomarkers). As the kidney donors remained anonymous, only age and sex were reported for the control group. Patient characteristics (A) Control: 55.9 ± 12.3 years, 30.77% male, renal dysfunction: N/A, (B) AAA: 66.5 ± 7.3 years, 54,6% male, 27.3% renal dysfunction (patient characteristics of 1 AAA patient unavailable). A list of included VSMC lines and the lines used in each experiment are shown in Table T1 and T2.

VSMC were isolated from aortic biopsies as described previously (19). Briefly, the adventitial layer and the intimal layer were removed from the biopsy, leaving a homogeneous and compact medial layer. This medial layer was sliced into approximately 8 pieces, which were placed in a T-25 flask filled with 1.5 ml culture medium. VSMC were cultured in a humidified incubator at 37 °C and 5% CO₂, in Human Vascular Smooth Muscle Cell Basal Medium (Gibco, Thermo Fisher Scientific) supplemented with Smooth Muscle Growth Supplement (Gibco, Thermo Fisher Scientific), 100 units/ml penicillin and 100 µg/ml streptomycin. Culture medium was refreshed once a week, cells split at 80-90% confluency. Primary VSMC were used between passages 1-8 in all experiments.

### hiPSC line culture and maintenance

The hiPSC line SCVI-111 used in the present study stems from the Stanford Cardiovascular Institute biobank (RRID:CVCL_C6U9). SCVI111 was Sendai virus reprogrammed from 490 peripheral blood mononuclear cells, the donor was a healthy male with normal karyotype. hiPSCs were maintained on Vitronectin (StemCell Technologies) coated suspension plates (Greiner) in mTeSR Plus (StemCell Technologies). Cells where passaged as colonies using Gentle Cell Dissociation Reagent according to the manufacturers instruction (StemCell Technologies), media change was performed daily.

### Differentiation of hiPSC-ECs

hiPSCs were directed toward an EC lineage following previously established protocols (20–22) with minor modifications. For the mesoderm induction phase (day 0–3), mTeSR Plus was substituted with B(P)EL medium containing 8 μM CHIR99021 (Tocris Bioscience). From day 3 onward, vascular specification was promoted by switching to B(P)EL supplemented with VEGF (50 ng/ml, PeproTech) and SB431542 (10 μM; Tocris Bioscience, 1614), with medium changes performed on days 3, 6, and 9. On day 10, ECs were selectively isolated using CD31- Dynabeads (Thermo Fisher Scientific), as previously described (20, 21). The purified hiPSC-derived ECs were expanded in Human Endothelial-Serum Free Medium (EC-SFM, Gibco) supplemented with 1% human platelet-poor serum (P2918, Sigma), VEGF (30 ng/ml, Peprotech), and bFGF (20 ng/ml, Miltenyi Biotec). Cells were cryopreserved at passage 1 in cryopreservation media (40% ECGM-2 (Promocell), 50% FBS (Gibco), 10% DMSO (Sigma Aldrich)).

### Immunocytochemistry 2D

Cells were seeded onto ibidi µ-Slide 8 Well slides (ibiTreat, ibidi GmbH, Germany), pre-coated with 0.1% gelatin. Fixation was carried out using 4% paraformaldehyde (PFA, ThermoFisher) in PBS(-/-) (Gibco) for 15 minutes at room temperature (RT). Following fixation, cells were washed three times with PBS(-/-) and permeabilized with 0.2% Triton X-100 in PBS(-/-) for 3 minutes. Cells were blocked in 1% human serum albumin in PBS(-/-) for 1 hour. Primary antibodies were applied in the same blocking buffer, incubated for 1 hour at RT (Antibody list Tab. T3) and washed three times with PBS(-/-). Cells were then incubated with the appropriate secondary antibody and fluorescent probes (Antibody list Tab. T3) for 1 hour at RT, following 3 wash steps with PBS(-/-).

### Setting up microvascular models on chip

C-VoCs and AAA-VoCs were generated as previously described with minor adjustments (18). Briefly, commercially available microfluidic devices (IdentX9, AIM Biotech) were employed. For microvascular network formation, primary C-VSMCs or AAA-VSMCs were combined with hiPSC-ECs to achieve a cell master mix suspension containing 10 × 10⁶ hiPSC-ECs/mL and 2 × 10⁶ VSMCs/mL, resulting in a 5:1 EC-to-VSMC ratio. The cell suspension master mix was prepared in ECGM-2 medium supplemented with VEGF (50 ng/mL) and thrombin (4 U/mL, Sigma Aldrich). Prior to loading, one part of cell suspension master mix was combined with an equal volume of ECGM-2 containing VEGF (50 ng/mL) and thrombin (4U/mL), followed by a 1:1 dilution with fibrinogen (Sigma-Aldrich) to reach a final fibrinogen concentration of 3 mg/mL. Immediately after, 15 µL of the resulting cell-hydrogel mixture was added into the central channel of the IdentX9 chip and allowed to polymerize for 15 minutes at RT. To establish gravity-driven perfusion, 100 µL of VEGF-supplemented ECGM-2 was added to the right media inlet and 50 µL to the left. Media was refreshed daily using the same approach, maintaining gravity-driven flow for 7 days. On the first day of culture, the medium was supplemented with the γ-secretase inhibitor DAPT (10 μM, Tocris Bioscience).

### Immunocytochemistry VoC

Microvascular networks were fixed directly within their microfluidic chips using 4% PFA for 30 minutes at RT. To permeabilize cellular membranes, samples were treated with 0.5% Triton X-100 for 15 minutes under the same conditions. Each of these steps was followed by three 10 minutes washes with PBS(-/-). Samples were blocked in PBS(-/-) containing 2% bovine serum albumin (BSA, Sigma Aldrich) for 3 hours at RT. Primary antibodies, diluted 1:400 in 1% BSA, were applied and incubated overnight at 4 °C (Antibody list Tab. T3). Following incubation, samples underwent three 15 minute washes with PBS(-/-) before exposure to secondary antibodies, diluted 1:600 in 1% BSA, for 2 hours at RT. Imaging was carried out on a Nikon AXR confocal microscope (Nikon Instruments Inc., Tokyo, Japan). Depending on the analysis performed, image processing and three-dimensional reconstructions were generated using Imaris (version 10.2.0, Bitplane, Oxford Instruments) or NIS-Elements (version 5.42.04, Nikon Instruments Inc., Tokyo, Japan).

### Permeability assessment of microvascular networks

To assess endothelial barrier function, a permeability assay was carried out. To visualize vascular structures, VoCs were incubated with Ulex Europaeus Agglutinin I (UEA-I, DyLight 649, DL-1068-1, Vector Laboratories; dilution 1:600) for 1 hour under standard incubation conditions (37°C, 5% CO₂). Live-cell confocal imaging was then performed using a Nikon AXR microscope equipped with environmental control. For the permeability test, 70-kDa Fluorescein Isothiocyanate-Dextran (FITC-Dextran, Merck; 2 µg/mL) prepared in ECGM-2 medium was added to the top left (100 µL) and right (50 µL) inlets of the microfluidic device. Fluorescence images were captured every 10 minutes for a total of 50 minutes. Quantification was conducted using NIS-Elements software. Briefly, regions of interest (ROIs) were determined by segmenting UEA-I–positive vascular areas and UEA-I–negative ECM regions. Mean fluorescence intensity (MFI) of FITC-Dextran within these ROIs was measured over time to evaluate tracer diffusion into the ECM, and a leakage dye index (LI) was calculated as the fraction of MFI in the vessel over MFI of ECM per timepoint.

### Geometry analysis of microvascular networks

On day 7 of culture, vascular networks were fixed and subjected to immunostaining using either UEA-I or PECAM1 to label endothelial structures. Overview images of microvascular networks were generated and subsequently transformed to as maximum intensity projection in Z. Image segmentation was carried out in CellProfiler software (version 4.2.1, Broad Institute). The resulting binary images were analyzed in ImageJ (NIH, USA) using the open-source DiameterJ plugin (23). All image processing and quantitative analyses followed identical workflows to ensure consistency across experimental groups.

### Shear stress application on hiPSC-ECs in 2D

A computer-controlled pump system (ibidi), consisting of a pump, fluidic unit, and perfusion set (15 cm tubing, 1.6 mm inner diameter, 10 ml reservoirs), was employed to culture hiPSC-ECs under laminar flow. Cells were seeded into fibronectin (Merck) coated μ-slides VI 0.6 (ibidi) at a density of 0.5 × 10⁶ cells/ml. After a 24-hour attachment period, the slides were connected to the perfusion system, and flow was applied gradually, starting from 1 hour at 2.5 dyn/cm², then 1 hour at 7.5 dyn/cm² and subsequently maintained at 18 dyn/cm² for three days. All flow experiments were performed in a standard cell culture incubator (37 °C, 5% CO₂), in EC-SFM supplemented with 1% human platelet-poor serum, VEGF (30 ng/ml) and bFGF (20 ng/ml).

### Determination of polarity index (PI)

EC alignment was assessed by measuring the orientation of the Golgi apparatus relative to the cell nucleus, from which a polarity index (PI) was calculated. The PI was defined as the mean resultant length of angles, transformed to center around the vertical axis using the formula: (𝜽 + 𝟗𝟎°) 𝒎𝒐𝒅 𝟏𝟖𝟎° – 𝟗𝟎°. Golgi structures were visualized using Golgin97 antibody (Antibody list Tab. T3). ECs were labeled with VE-cadherin (Antibody list Tab. T3), and nuclei with DAPI (Dako, 1:600). A parallel grid was drawn relative to the top and bottom image borders to establish a reference axis (Fig. S1). Using the angle measurement tool, vectors were drawn from the center of the nucleus to the center of the Golgi-apparatus, with the second axis aligned parallel to the grid lines. The resulting angles were used to determine the Golgi-to-nucleus orientation per cell. For microvascular networks, the microvascular structures were visualized with UAE-I and Golgi-apparatus and nucleus as described above. The reference grid was determined parallel to the vessel walls (Fig. S1). Additionally, VoC images were subdivided into straight vessel segments (areas without bifurcations) and branching regions (areas with two or more vessel branches). Images were analyzed using ImageJ.

### Calcium transient determination

Two hours prior to the assay, culture medium was replaced with a mixture consisting of (A) ECGM-2 medium supplemented with VEGF (50 ng/mL) and UEA-I, and (B) 5 µM Cal-520 (Abcam) in 0.02% Pluronic F-127 (Sigma Aldrich). Live-cell imaging was conducted using a Nikon AXR confocal microscope equipped with environmental control (37 °C, 5% CO₂). Baseline calcium activity was recorded for 2 minutes at 3-second intervals. Endothelin-1 (ET- 1) was then introduced into the media inlets of the microfluidic platform to achieve a final concentration of 10 nM. After a 5-minute incubation period, the same imaging coordinates were revisited and calcium dynamics were recorded for an additional 2 minutes at the same acquisition interval. ROIs corresponding to VSMCs were identified based on peak calcium loading following ET-1 stimulation (Fig. S2). Quantification was performed on z-stack images, and the MFI of Cal-520 was measured at each time point for individual ROIs to assess temporal calcium fluctuations.

### Cytokine detection array

Proteome Profiler Human Cytokine Array Kit (R&D Systems) was used to quantify the cytokine levels in VoC supernatants. Supernatants were collected from three VoCs (Control, AAA) 24 hours after the final media exchange and pooled before processing. The arrays were performed following the instructions provided by the manufacturer with one modification where SuperSignal™ West Femto Maximum Sensitivity Substrate from Thermo Fisher Scientific was applied for the detection and imaging of the resulting dot blot signals.

### Statistical analysis

All functional assays on VoCs were performed on culture day 7. Statistical analysis were performed using GraphPad Prism version 10 (GraphPad Software, San Diego, CA). Data are presented as mean ± SEM or mean ± SD. Data were tested for normality, and appropriate parametric or non-parametric tests were applied accordingly. For longitudinal data, repeated-measures analyses were used where appropriate. Statistical significance was set at *p* < 0.05 (\**p* < 0.05, \*\**p* < 0.01, \*\*\**p* < 0.005, \*\*\*\**p* < 0.001).

## Results

### C-VSMCs and AAA-VSMCs support hiPSC-EC vascular network formation in 3D

In this study, we generated a 3D *in vitro* co-culture model of the microvasculature in the AAA aortic wall using a healthy donor derived hiPSC-EC line in co-culture with primary C-VSMCs or primary AAA-VSMCs (Fig. 1A). In 2D *in vitro* culture, C-VSMCs and AAA-VSMCs express canonical VSMC markers including smooth muscle protein 22-alpha (SM22α) and alpha-smooth muscle actin (αSMA) (Fig. 1B). hiPSC-ECs form a confluent monolayer and express key endothelial markers, including vascular endothelial cadherin (VE-cadherin) and Platelet Endothelial Cell Adhesion Molecule-1 (PECAM-1/CD31) (Fig. 1B).

We found that co-culture of hiPSC-ECs with C-VSMCs and AAA-VSMCs resulted in vascular networks after 7 days (Fig. 1C). For both C-VoCs and AAA-VoCs, we observed vascularization of the entire hydrogel chamber in the x-y axis (Fig. 1C). We furthermore observed that both C- VSMCs and AAA-VSMCs enabled vascular lumen formation, as evidenced by open microvessels visible in the central planes of z-stack imaging (Fig. 1C) and y-z orthogonal views of vascular lumen with lumenized microvascular structures for C-VoCs and AAA-VoCs (Fig. 1C). VSMCs remained viable for 7 days of co-culture, retained their characteristic phenotype as indicated by SM22 staining, and were uniformly distributed throughout the hydrogel channel. We furthermore show that hiPSC-EC identity is preserved in co-culture with C- VSMCs and AAA-VSMCs. Microvascular structures showed organized adherens junctions (AJ) both in C-VoCs and AAA-VoCs structures, identified by VE-Cadherin and PECAM-1 expression (Fig. 1D) and tight junction protein Claudin – 5 (CLDN-5, Fig. S3) on the entire network area. Moreover, we assessed whether hiPSC-ECs and VSMCs establish heterotypic cell-cell contact within the co-culture. To do so, cells were stained with a DNA dye and phalloidin to visualize nuclei and F-actin, respectively. Vascular structures were identified by VE-cadherin and PECAM-1 staining, allowing VSMCs to be distinguished as phalloidin/DNA-positive but endothelial marker-negative (Fig. 1E). After 7 days of co-culture, we observed multiple contact points between VSMCs and hiPSC- ECs (Fig. 1E). This was found throughout the entire hydrogel channel and was visibly comparable for C-VoCs and AAA-VoCs, indicating that in both systems, VSMCs are spatially interacting with the microvascular network.

### AAA-VoCs show enlarged vascular diameter

We next aimed at characterizing the geometry of the microvascular structures generated by hiPSC-ECs in co-culture with either C-VSMCs or AAA-VMCs. Vascular structures were identified by PECAM-1 staining and analysed as z-stack projections (Schematic overview of workflow: Fig. 2A). Morphometric analysis showed that the hiPSC-ECs in co-culture with AAA- VSMCs generated microvascular structures with increased average vessel diameters (Fig. 2B). We furthermore found that the maximum vessel diameter was also increased in AAA- VoCs compared to C-VoCs (Fig. 2C). Moreover, we quantified branching points and length of vasculature per channel. For these two parameters, we did not find a significant difference between C-VoCs and AAA-VoCs (Fig. 2D, Fig. 2E). We furthermore investigated whether the increased vessel diameter in AAA-VoCs correlates with an increased hiPSC-EC proliferation. We therefore stained day 7 VoC-cultures for Ki67, a nuclear marker for cellular proliferation, along with a DNA dye, and identified hiPSC-ECs using an additional UAE-1 stain, subsequently determining the percentage of Ki67-positive nuclei among hiPSC-ECs (Fig. S4). We found a significantly increased fraction of Ki67 positive hiPSC-EC nuclei in AAA-VoCs over C-VoCs (Fig. 2F), showing that the increased vascular diameter in AAA-VoCs correlates with an increased hiPSC-EC proliferation. Overall, these results indicate that both C-VSMCs and AAA- VSMCs support vascular network formation of comparable complexity, but that the vascular diameter is increased when hiPSC-ECs are co-cultured with AAA-VSMCs as compared to C- VSMCs, which is correlated with increased hiPSC-EC proliferation.

**Figure 2.**
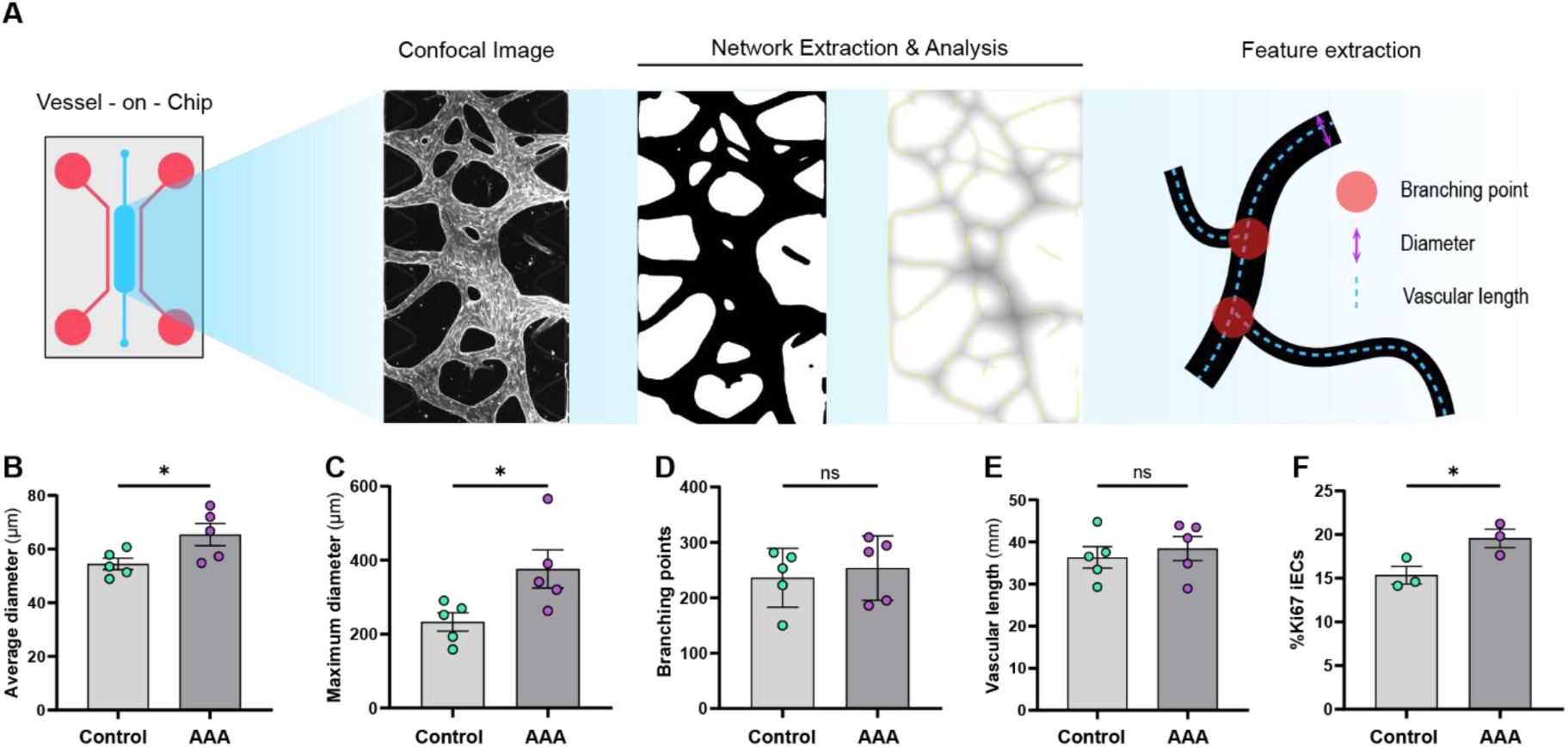
Geometry of microvasculature in VoC models. (A) Schematic overview of workflow for geometry analysis in VoC models. (B – E): Quantification of microvascular geometry in VoC models. (B) microvascular diameter in µm, (C) maximal vascular diameter in µm, (D) number of branching points (E) total vascular length in µm. Data are shown as mean ± SEM. Data points represent 5 independent experiments on 5 (control) and 4 (AAA) independent VSMC lines; 2–5 VoC channels per experiment were imaged and data averaged to yield one value per cell line. Data passed Shapiro–Wilk normality test, groups compared using unpaired two-tailed t-test. *p< 0.05. (F) Quantification of fraction of Ki67 positive hiPSC-ECs nuclei at day 7 in %. Data are shown as mean ± SEM. Data points represent 3 independent experiments on VoCs with 3 (control) and 3 (AAA) independent VSMC lines; 3–6 VoC channels per experiment were imaged and data averaged to yield one value per cell line. Data passed Shapiro–Wilk normality test, groups compared using unpaired two-tailed t-test. *p< 0.05.

### hiPSC-ECs align with gravity driven flow in both C-VoCs and AAA-VoCs

A hallmark of viable ECs is their alignment under laminar shear stress (LSS), and the reduction of the alignment in areas of turbulent flow, which is atheroprotective and mediated by intrinsic mechanosensitivity (24). However, flow alignment is also linked to crosstalk with the microenvironment of the aortic wall, primarily composed of VSMCs (25, 26). Therefore, a lack of hiPSC-EC polarization under flow may reflect not only impaired endothelial mechanoresponsiveness but could also be a consequence of a dysregulated or non-physiological phenotype of the co-cultured VSMCs.

To assess flow responsiveness, we utilized the well-established phenomenon that under LSS, ECs polarize, with the Golgi aligning upstream of the nucleus (Fig. 3A) (27). The polarity index (PI) was defined as the combined degree of the angles transformed to center around the vertical axis. Values close to 1 indicate strong directional alignment, while values near 0 reflect random or dispersed orientations. Next to VoC models (Fig. 3B), we included 2 control groups: hiPSC-ECs in a 2D *in vitro* static culture (without flow) (Fig. 3B) and hiPSC-ECs in a 2D *in vitro* setup under defined LSS (72 hours, 18 dyn/cm²). Nuclei, Golgi apparatus and ECs were visualized using respective fluorescent labels in 2D cultures after 72h; and in VoC cultures after 7 days (Fig. 3B). In VoC cultures, we furthermore separated straight vascular areas (areas of the vasculature without branching vessels, Fig. 3B) from branching points (areas of the vasculature with 2 or more branching vessels, Fig. 3B), to investigate whether flow alignment differs between these two areas, indicative of laminar and turbulent flow. The direction of flow (DOF) in VoCs was determined by estimating a parallel line along the vascular wall to be analyzed (Fig. 3B).

**Figure 3.**
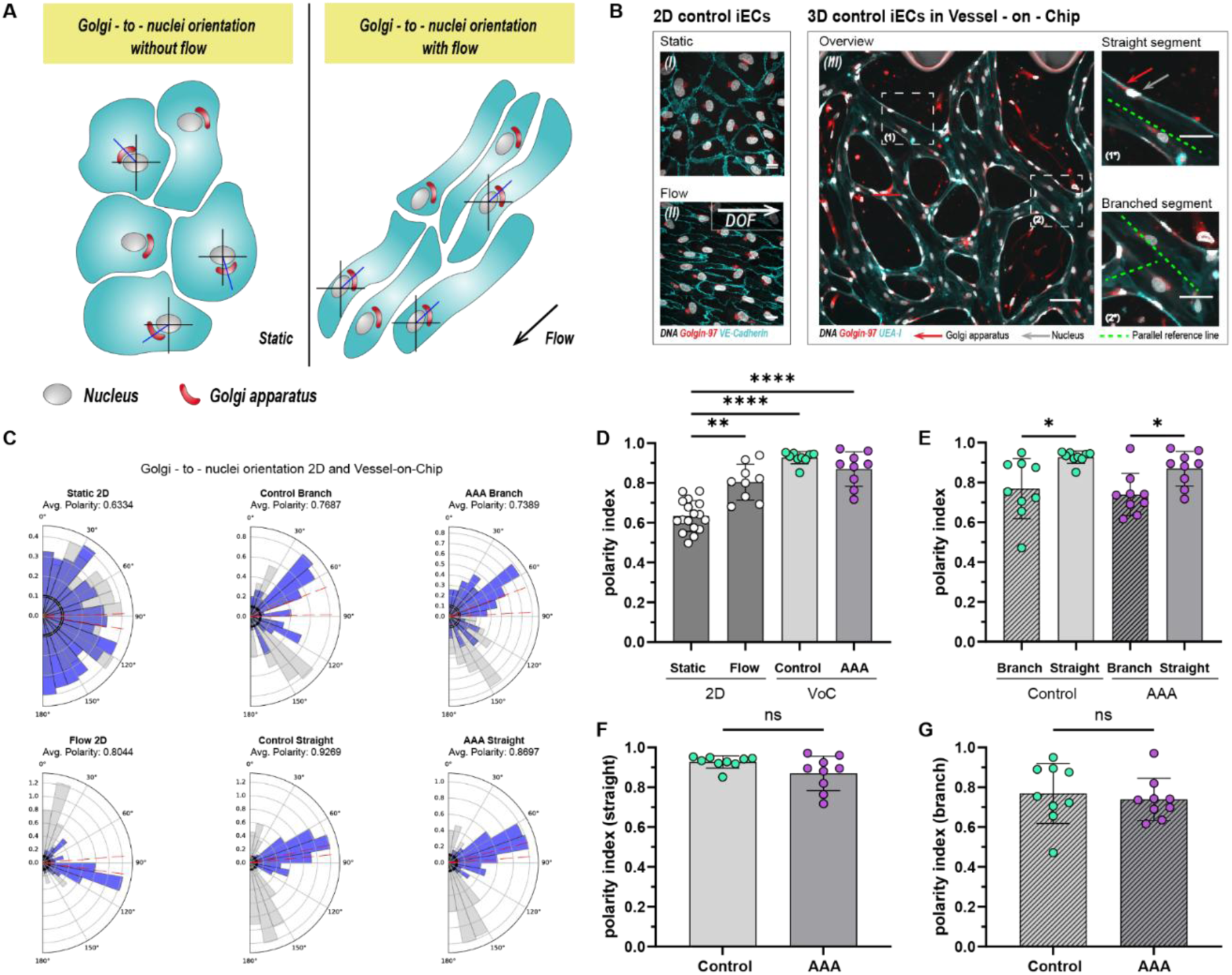
Endothelial polarity in the VoC models. (A) Schematic overview of EC Golgi-to-nuclei orientation in static conditions (left panel) and upon laminar shear stress exposure (right panel), as well as Golgi-to-nuclei angles (blue line) indicating the angle used to calculate the polarity index (PI). (B) Representative confocal images of hiPSC-ECs in 2D for Golgi-to-nuclei orientation analysis (left panel, (I) static, (II) flow), (gray: DAPI, red: Golgin – 97, cyan: VE-Cadherin). Scale bars: 20 µm. DOF = Direction of flow in Ibidi flow set-up. (III) Representative confocal image of VoC for Golgi-to-nuclei orientation analysis. Scale bars: 100 µm. (1*) Images of a representative straight labelled segment and (2*) branched segment derived from indicated areas (1) and (2) respectively, with green dashed line indicating the determined reference line for Golgi-to-nuclei angle determination (gray: DAPI, red: Golgin – 97, cyan: VE-Cadherin). Scale bars: 50 µm. (C) Radial histograms depicting the distribution of Golgi-to-nuclei orientation in hiPSC-ECs for static and flow 2D (first column), C-VoC branched and straight areas of the microvasculature (middle column) and AAA-VoC branched and straight areas of the microvasculature (right column). Blue: transformed angles, gray: original angles, red dashed lines indicating the lower and upper limits of the 95% confidence interval of the transformed orientation angles. Radial axis indicates density (frequency) of observations normalized to unit area. (D-G) Quantification of Golgi-to-nuclei orientation. (D) PI comparison in 2D static and flow, and straight branches of C-VoC and AAA-VoC, (E) PI in branched and straight areas of C-VoC and AAA-VoC. (F) PI for straight areas of C-VoC and AAA-VoC, (G) PI for branched areas of C-VoC and AAA-VoC. (D) Data are shown as mean ± SD. Data points in graph represent for Static 2D: 3 independent experiments (5 - 6 fields of view); for Flow 2D: 3 independent experiments (3 fields of view each); for C- and AAA-VoC: 3 independent experiments with 3 independent VSMC lines (3 fields of view each). (E-G) Data are shown as mean ± SD. Data points in graph represent 3 experiments with 3 independent VSMC lines (3 fields of view each). C-VoC (straight) failed normality (Shapiro-Wilk test, W = 0.7673, p = 0.0086). Pairwise comparisons performed using unpaired two-tailed t-tests for normally distributed groups and Mann–Whitney tests for non-normal groups; ****p< 0.001, *** p < 0.005, ** p< 0.01, * p< 0.05.

PI quantification for each condition is visualized in radial histograms (Fig. 3C). We found that for static conditions, the PI of hiPSC-EC is comparably low and that the application of flow in 2D for 72h significantly increases the PI (Fig. 3D), showing that these hiPSC-EC are flow- responsive. Interestingly, we found that for straight branches in both C-VoCs and AAA-VoCs, the PI was further increased when compared to the 2D flow condition (Fig. 3D). We furthermore found that there is a significant difference between branched and straight areas of the vascular networks in C-VoCs and AAA-VoCs, where straight areas show an increased PI (Fig. 3E). When comparing C-VoC PIs with AAA-VoC PIs for straight areas (Fig. 3F) or branched areas (Fig. 3G), we did not find a significant difference.

In sum, these findings strengthen our findings that both C-VSMCs and AAA-VSMCs support hiPSC-ECs successfully in a microvascular model and that hiPSC-ECs are viable, showing key physiological features such as alignment in the direction of flow.

### AAA-VSMCs show increased cell number and surface area at day 7 of AAA-VoC culture

Having established that both C-VSMCs and AAA-VSMCs are viable and able to support a microvascular network *in vitro*, we sought to examine whether AAA-VSMCs recapitulate AAA specific features in our model. We employed immunostaining using PECAM-1 to label microvascular structures and SM22 to identify VSMCs. Three-dimensional rendering and quantitative analysis were subsequently performed using to assess the spatial organization, count and size of C-VSMCs and AAA-VSMCs after 7 days on-chip (Fig. 4A). A hallmark of AAA is the so called VSMC phenotypic switch from a quiescent, contractile phenotype towards a proliferative and synthetic phenotype (28) and previous work identified that proliferative genes were significantly increased in human AAA tissue (29). Interestingly, we found an increased number of AAA-VSMCs after 7 days in VoC culture compared to C-VSMCs (Fig. 4B). Quantitative morphometric data on VSMC dimensions in human AAAs are currently lacking. However, phenotypic switching from spindle-shaped contractile cells toward irregular synthetic, fibroblast- or macrophage-like morphologies in AAA-VSMCs suggest a change in cell morphology. We found in our system that AAA-VSMCs show an increased surface area compared to C-VSMCs after 7 days of hiPSC-EC co-culture, supporting that hypothesis and further suggesting the presence of a preserved disease phenotype of AAA-VSMCs (Fig. 4C). We furthermore investigated whether AAA-VSMCs are aberrant in terms of distance to the microvascular network (Fig. 4D) or amount of area covered on the microvascular network (Fig. 4E) and did not find a significant difference between AAA-VoCs and C-VoCs. In sum, we find that AAA-VSMCs have structural and behavioral differences on-chip that is reflected by increased VSMC number after 7 days in culture and an distinct morphology, but their behavior in relation to the microvascular network is comparable to C-VSMCs.

**Figure 4.**
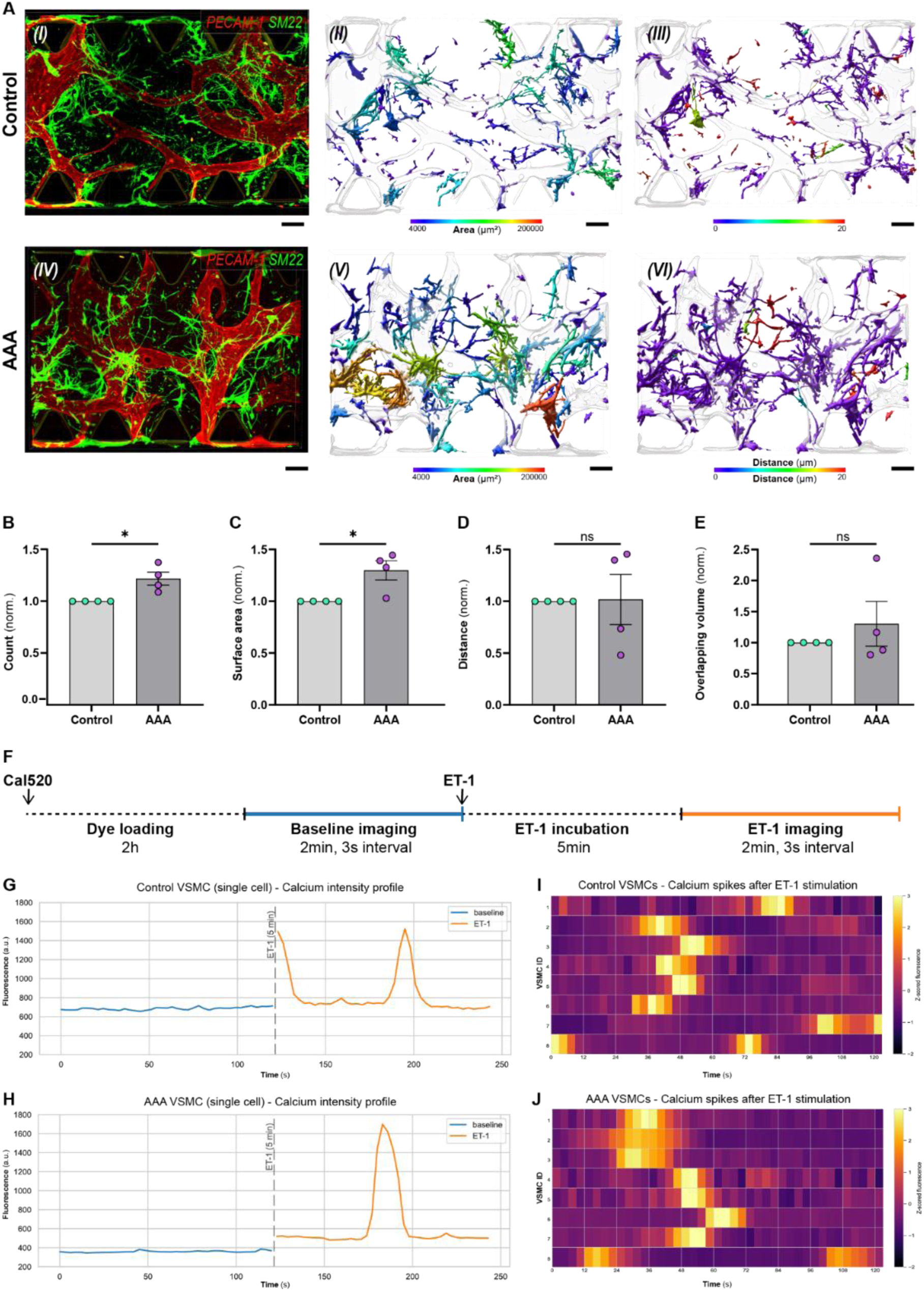
AAA-VoC VSMC show increased cell number but no increase in distance to ECs or Ca^2+^-responses. (A) Representative confocal images of (I) C-VoC and (IV) AAA-VoC showing hiPSC-ECs (red: PECAM-1) and C-VSMCs (green: SM22). Surface-rendered objects based on PECAM-1 (microvascular network, transparent render) and SM22 (VSMCs, color-coded render). (II + V) VSMCs with a color-coded scale for 3D surface area in µm² and (III + VI) color-coded scale for VSMC distance to microvascular network in µm. (B-E) Quantification of VSMC characteristics in C-VoC and AAA-VoC showing (B) normalized cell count, (C) normalized surface area, (D) normalized VSMC distance to microvascular network and (E) normalized overlapping volume between VSMCs and microvascular network.Data are shown as mean ± SEM. Data points represent n = 3 independent VSMC lines (control and AAA), each assayed in 4 separate experiments; 3–6 VoC channels per experiment were imaged and data averaged to yield one value per cell line. AAA-VoC data points were normalized to the corresponding C-VoC processed in parallel. Unnormalized data passed Shapiro–Wilk normality test. Unpaired two-tailed t-test. *p = < 0.05. (F) Schematic overview of experimental timeline for calcium transient recordings. (G) Representative graph for ET-1 responsive C- VSMC, blue line indicating calcium activity at baseline before ET-1 stimulation, orange line after ET-1 stimulation. Dashed line indicating incubation step of 5 minutes after ET- 1 stimulation, in between baseline and ET-1 recording. (H) Representative graph for ET-1 responsive AAA-VSMC, blue line indicating calcium activity at baseline before ET-1 stimulation, orange line after ET-1 stimulation. Dashed line indicating incubation step of 5 minutes after ET-1 stimulation, in between baseline and ET-1 recording. (I) Heatmap with a z-score normalized color-code for calcium activity depicting 8 representative C-VSMCs (VSMC ID 1-8) 5 minutes after ET-1 stimulation. (J) Heatmap with a z-score normalized color-code for calcium activity depicting 8 representative AAA-VSMCs (VSMC ID 1-8) 5 minutes after ET-1 stimulation.

### C-VSMCs and AAA-VSMCs respond to ET-1 after 7 days of co-culture on-chip

To further characterize VSMC dynamics in our system, we investigated whether C-VSMCs and AAA-VSMCs exhibit calcium transients in response to a contractile stimulus (30). We employed the fluorescent calcium indicator Cal-520 AM for live-cell imaging (31, 32). We imaged for 2 minutes at baseline, and for 2 minutes after stimulation with ET-1 (schematic experimental overview: Fig. 4F), a vasoconstrictive peptide primarily produced by ECs and known to trigger calcium signaling in VSMCs *in vivo* (33). Fields of view were selected based on UEA-I staining, indicating vascularized hydrogel areas, and ROIs were selected for calcium- dye positive, UEA-I negative cells, representing VSMCs (Fig. S2). We then plotted the MFI for single ROIs before; and 5 minutes after ET-1 stimulation and found ET-1 responsive cells both in C-VoCs (Fig. 4G) and AAA-VoCs (Fig. 4H). Furthermore, we found in both models comparable calcium transient frequency and duration for VSMCs, indicating that both C-VoCs and AAA-VoCs contain viable VSMCs with the ability to respond in a similar manner to physiological stimuli such as ET-1 (Fig. 4I and 4J).

Together, these results indicate that although AAA-VSMCs have a distinct phenotype in VoCs, viability and functionality as deduced by calcium transient remains comparable between C- VSMCs and AAA-VSMCs.

### AAA-VoCs show an elevated level of pro-inflammatory cytokine expression

We lastly evaluated whether our *in vitro* models reflect AAA-specific features, including endothelial dysfunction (34) and increased inflammatory cytokine levels (35). We tested whether the pro-inflammatory cytokine expression profile differs between C-VoCs and AAA- VoCs. To that end, we performed a human cytokine array assay on VoC media supernatant that was collected at day 7, after 24 hours of incubation on VoCs (Fig. 5A). For AAA-VoC supernatant, we observed elevated levels of CXCL1, CXCL12, CCL2, IL-8, PAI-1, G-CSF and MIF compared to C-VoC supernatant (Fig. 5B, C). To further dissect the extent of cytokine elevation in our system we plotted a volcano plot for detected cytokines (Fig. 5D), revealing that although all cytokines (except IL-6) show elevated levels in AAA-VoC, they are not significantly increased when observed individually.

**Figure 5.**
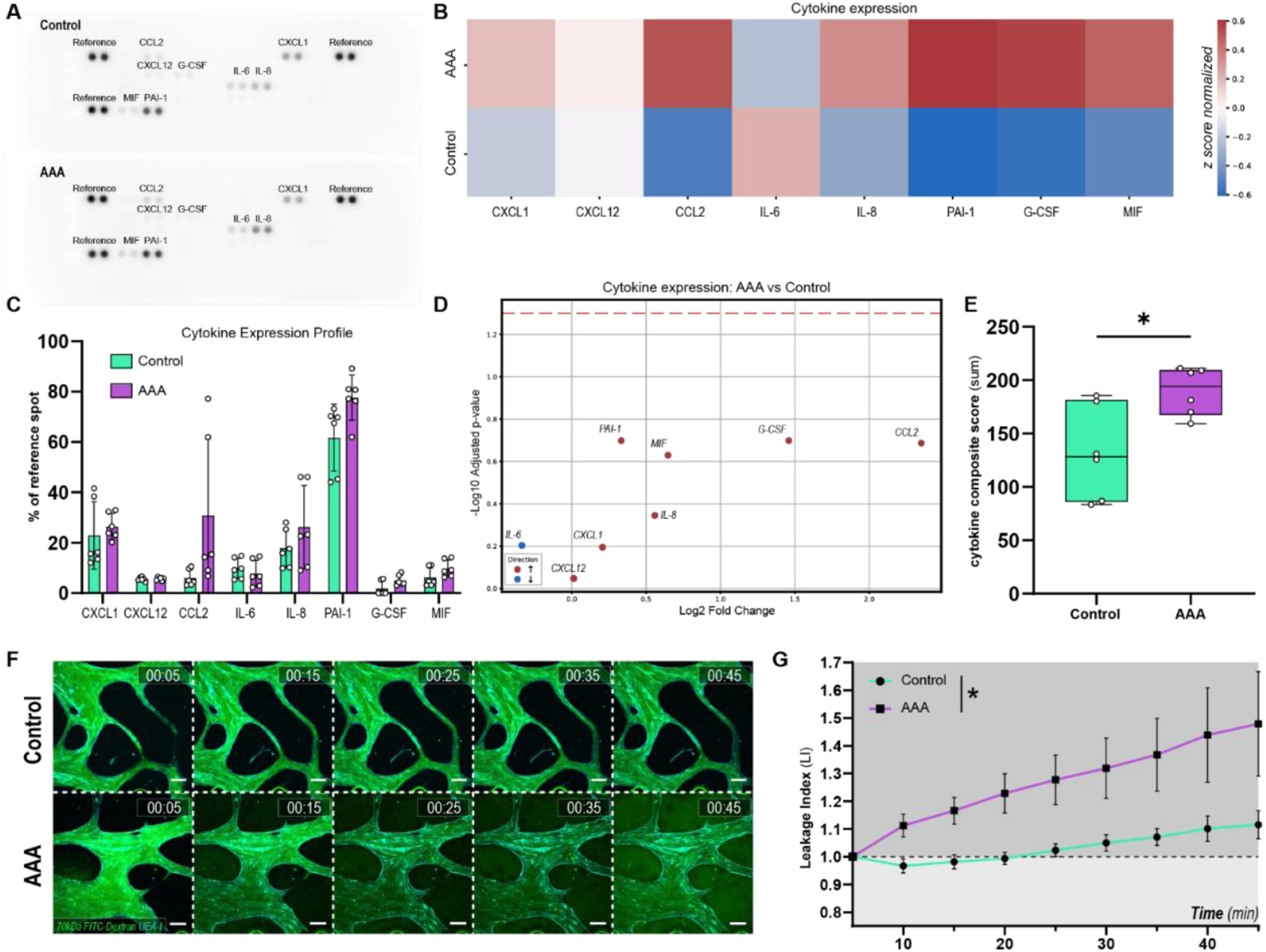
Cytokine expression and endothelial barrier in VoC models. (A) Representative human cytokine array detection membrane after incubation with C- VoC supernatant (top panel) or AAA-VoC supernatant (bottom panel). (B) Z-score normalized heatmap for cytokine expression levels in C-VoC and AAA-VoC supernatants, data from MFI quantification of cytokine array kit.(C) Expression levels of cytokines after MFI detection in cytokine array kit shown as % MFI of reference spot. Data are shown as mean ± SD, from n = 3 independent VSMC lines (control and AAA), each assayed in 3 separate experiments, 2 technical replicates per experiment. (D) Volcano plot showing log₂ fold change versus –log₁₀ adjusted p-value for cytokine expression in AAA-VoC against C-VoC, with red and blue dots indicating up- or downregulated cytokines in AAA-VoC, and the red dashed line marking the significance threshold (adjusted p = 0.05). (E) Composite score of cytokine expression, calculated as the sum of expression levels for all measured cytokines compared between groups using an unpaired two-tailed t-test after confirming normality. *p < 0.05. Data are shown as box plots: median (line), interquartile range (box), and minimum to maximum values. Each point represents an individual sample (n = 3 independent VSMC lines, 2 technical replicates each). (F) Representative images of microvasculature (cyan: UEA-I) perfused with 70 kDa FITC-Dextran (green) over the time course of 45 minutes. Top panel C-VoC, bottom panel AAA-VoC (Time stamp: HH:MM). Scale bars: 100 µm. (G) Quantification of FITC–dextran leakage across the microvascular barrier in C-VoCs and AAA-VoCs, expressed as the ratio of mean MFI in the ECM to MFI in the vasculature over time. Data are shown as mean ± SEM, from 4 (control) and 5 (AAA) independent experiments. 2 – 6 VoC channels imaged per experiment. Tested for main effect of group across all timepoints, two-way repeated measures ANOVA. *p < 0.05.

Since no individual cytokine showed a significant difference between C-VoCs and AAA-VoCs, we next asked whether the combined cytokine response might uncover a grouped difference in cytokine expression that is not apparent on a single cytokine level. To this end, we calculated a composite score by summing the concentration of all measured cytokines for either C-VoCs or AAA-VoCs, reflecting total cytokine burden. This composite score was significantly higher in AAA-VoCs compared to C-VoCs (Fig. 5E). Because cytokines differ in absolute scales, we also derived an alternative composite score by first log-transforming and then z-score normalizing each cytokine. This scale independent composite score showed a similar trend toward higher values in AAA-VoCs, although it did not reach statistical significance (*p* = 0.0626; Fig. S5).

Altogether, these results point out that AAA-VoCs show a moderate increase in cytokine expression for a variety of cytokines tested, and that the total expression levels of cytokines is significantly elevated in AAA-VoCs.

### AAA-VoCs exhibit increased vascular leakage

Lastly, we investigated vascular barrier integrity in AAA-VoCs. Endothelial dysfunction is a key feature of AAA, and endothelial barrier promoting genes have been reported downregulated in human aortic AAA tissue (36). To that end, we performed live cell imaging on day 7 VoCs (Vid. S1, S2). Vascular structures were highlighted using UAE-I and microvascular lumen were perfused with fluorescein isothiocyanate–conjugated dextran (70 kDa FITC-dextran) (Fig. 5F). In line with our initial findings that microvascular structures are lumenized, we confirmed that both C-VoCs and AAA-VoCs are perfusable. However, when quantifying vascular permeability, we found that AAA-VoCs show an increased vascular leakage over time as compared to C- VoCs (Fig. 5G). These results reveal that AAA-VoCs recapitulate a further aspect of AAA *in vitro*, namely endothelial dysfunction, reflected by an impaired endothelial barrier integrity. In combination with the elevated cytokine levels of AAA-VoCs described above, these findings suggest that AAA-VSMCs maintain key aspects of AAA, that manifest in 3D VoC models *in vitro*.

## Discussion

In the present study, we describe the development of a 3D self-organized model for the microvasculature in AAA using a healthy hiPSC-EC line to generate microvascular networks in co-culture with primary VSMCs from either healthy donors or AAA patients. With this model, we sought to address the current challenges in developing therapeutics for AAA, which are largely constrained by the limited translatability of animal studies and the vulnerability of the patient population.

To the best of our knowledge, we describe the first self-organizing 3D model of the aortic microcirculation in AAA. We present a scalable model on a commercial platform, allowing cells to self-organize into lumenized and perfusable microvascular networks within 7 days, thereby overcoming previous limitations.

To enable a robust comparison of the role of C- or AAA-VSMCs in the microvascular network, we decided to use a single hiPSC-derived EC line from a healthy donor in all models generated herein, resulting in an isogenic microvasculature, supported by either C- or AAA-VSMC lines.

We show that both primary C-VSMCs and AAA-VSMCs can be incorporated into a commercial microphysiological platform and support the formation of complex lumenized microvascular networks in co-culture with hiPSC-ECs. We show that VSMCs from both control and AAA donors are viable after 7 days of VoC culture. Both C-VSMCs and AAA-VSMCs were found to respond to ET-1 stimulation and hiPSC-ECs in both models align in the direction of flow. Importantly, we found that AAA-VSMCs preserve a disease-mimicking phenotype as reflected by an increase in cell number, microvessel surface area, cytokine production and microvascular permeability.

We found that AAA-VoCs are characterized by an increased average vascular diameter compared to C-VoCs. Previous studies showed that the average vascular diameter in self-organizing networks depends on ECs, supporting cell types, hydrogel composition, and the microfluidic platform used (37). On the same platform and fibrin hydrogel concentration as used in the current work, one study using hiPSC-ECs and human brain vascular pericytes (HBVPs) reported an average diameter of ∼53 μm, which was further reduced to ∼49–40 μm when astrocytes were introduced additionally (38). In the current model, hiPSC-ECs self-organized in co-culture with primary VSMCs into microvascular structures with an average vascular diameter of 54,50 µm (C-VoC) and 65,44 µm (AAA-VoC). Currently, there is limited understanding of how the number and type of mural support cells directly influence the self-organization and structural maturation of vascular networks. However, a common observation is that ECs in hydrogels alone form large, disorganized vessels, but adding mural cells yields finer, microvascular-like networks, underscoring their role in vessel patterning. In our model, while the number of AAA-VSMCs is increased, the resulting vessels are comparatively larger than those formed in the presence of C-VSMCs. This shows that AAA- VSMCs influence microvascular network formation on-chip, highlighting VSMC to EC cellular crosstalk in the present model and a distinct AAA-VSMC phenotype that differs from C-VSMCs.

We furthermore quantified EC polarization and compared 2D flow conditions with 3D VoC models. A potential limitation is the undefined shear rate in the 3D model compared to the precisely controlled 18 dynes/cm² achievable in the ibidi 2D system. A study using the same microfluidic platform and gravity driven flow as employed here, reported a calculated wall shear stress of 0.056-0,14 dynes/cm² (18). Although we did not investigate the shear stress values in our model, we estimate it to be in a comparable range, which is markedly lower than the 18 dynes/cm² we applied in our 2D ibidi flow control. It was therefore contrary to our expectations that flow alignment in the VoC, despite the lower flow rate compared to the ibidi system, was even more pronounced. The planar cell polarity (PCP) pathway is a non-canonical Wnt pathway that determines PCP in ECs (39). Although strongly activated by LSS (40), the PCP is also active during angiogenesis (41) and was suggested to play a role in vasculogenesis (42). It is likely that the strong polarity observed in our models arises from multiple PCP-activating cues resulting in an increased PI compared to the 2D culture where shear stress is the only PCP activator. The reduction of PI in branched areas of the VoC however indicates that shear stress plays a role in our system, as reduced or turbulent flow near bifurcations, correlates with a reduction of planar polarity (24).

We furthermore found that with an equal seeding density of VSMCs at day 0, AAA-VSMCs were higher in number at day 7 compared to C-VSMCs and showed an increased surface area. We tested for proliferation differences at day 7 of co-culture on-chip and did not find an increased number of Ki67 positive nuclei at this timepoint (Fig. S6). We furthermore tested whether AAA-VSMC proliferation or surface area differ in standard 2D cultures from C-VSMCs and found no significant difference (Fig. S7). Combined, these results indicate that the increased cell count and surface area found in our chips at day 7 is established initially after VSMC are embedded in hydrogel and co-cultured with hiPSC-ECs. This likely mimics a physiologically more relevant geometry than standard 2D cultures and thus better preserves the cellular characteristics. When comparing ET-1 induced calcium transients in C- and AAA- VoCs, we found comparable transients for VSMCs in both models. In AAA, VSMCs are known to shift towards a proliferative, less contractile phenotype (43), though the extent of contractile loss remains unclear. A recent *in vitro* study on AAA patient-derived VSMC lines reported reduced contraction in only 23% of cases, suggesting this phenomenon occurs but is relatively infrequent (19).

We lastly tested whether our model recapitulates AAA disease aspects *in vitro*, and found that, in line with *in vivo* observations, cytokine production is increased and endothelial barrier integrity is reduced for AAA-VoCs. The relative downregulation of IL-6 in AAA-VoCs was unexpected due to previous reports of IL-6 upregulation in rodent AAA models and patient samples (44, 45). This might be due to the broad activation spectrum of IL-6. It is known that VSMCs upregulate IL-6 in response to fibrin degradation products (46, 47) which could be a confounding factor after 7 day culture in fibrin hydrogel. However, we found an increased cytokine production in AAA-VoCs over C-VoCs in all other cytokines tested, with CCL2 showing the most pronounced differential expression for AAA-VoCs. CCL2 was found upregulated in a rodent model of AAA (48). Moreover, a recent study performing RNAseq on human AAA and control donor aortic tissue identified CCL2 as one of 6 key genes differentially expressed in AAA (49). We furthermore found that IL-8 is strongly upregulation in our model, and an upregulation of IL-8 in human AAA biopsies was reported previously (50). Altogether, we find in our model a cytokine expression profile for AAA-VoC that, to some extent, recapitulates findings from prior human patient studies. Furthermore, we show in the permeability assay that the microvasculature of AAA-VoCs shows a reduced barrier function, likely due to an interplay of the increased cytokine levels in AAA-VoCs and the established increased proliferation of hiPSC-ECs in AAA-VoCs.

In summary, we developed a model that recapitulates the microvasculature of the aortic wall *in vitro*, and by incorporating primary AAA patient derived VSMCs, we were able to mimic key aspects of the disease. Due to the commercial background of this platform, this model can be scaled up in a standardized way. The AAA-VoC model presented here provides a human-based platform to investigate and quantify alterations in AAA EC-VSMC crosstalk and cellular function. It holds the potential to overcome the current translational limitations of animal models and allows exploring novel therapeutic research avenues for improved understanding of the molecular regulation of AAA to ultimately develop efficient treatments for affected patients.

## Acknowledgments

The authors thank Prof. Joseph C. Wu from Stanford University for providing the hiPSC line SCVI-111. The authors furthermore thank Beau F. Neep of the Amsterdam UMC (Department of Respiratory Medicine) for material support, Victor L. H. Janssen of the Amsterdam UMC (Department of Vascular Surgery) for support with patient data management and Prof. Reinier A. Boon for carefully reading the manuscript and providing valuable input.

## Supplemental Material

Tables T1–T3 (they also called them S1-S3?) Figure 1 – 5

Supplementary figures S1 – S7 Videos S1–S2

---

**Figure S1:**
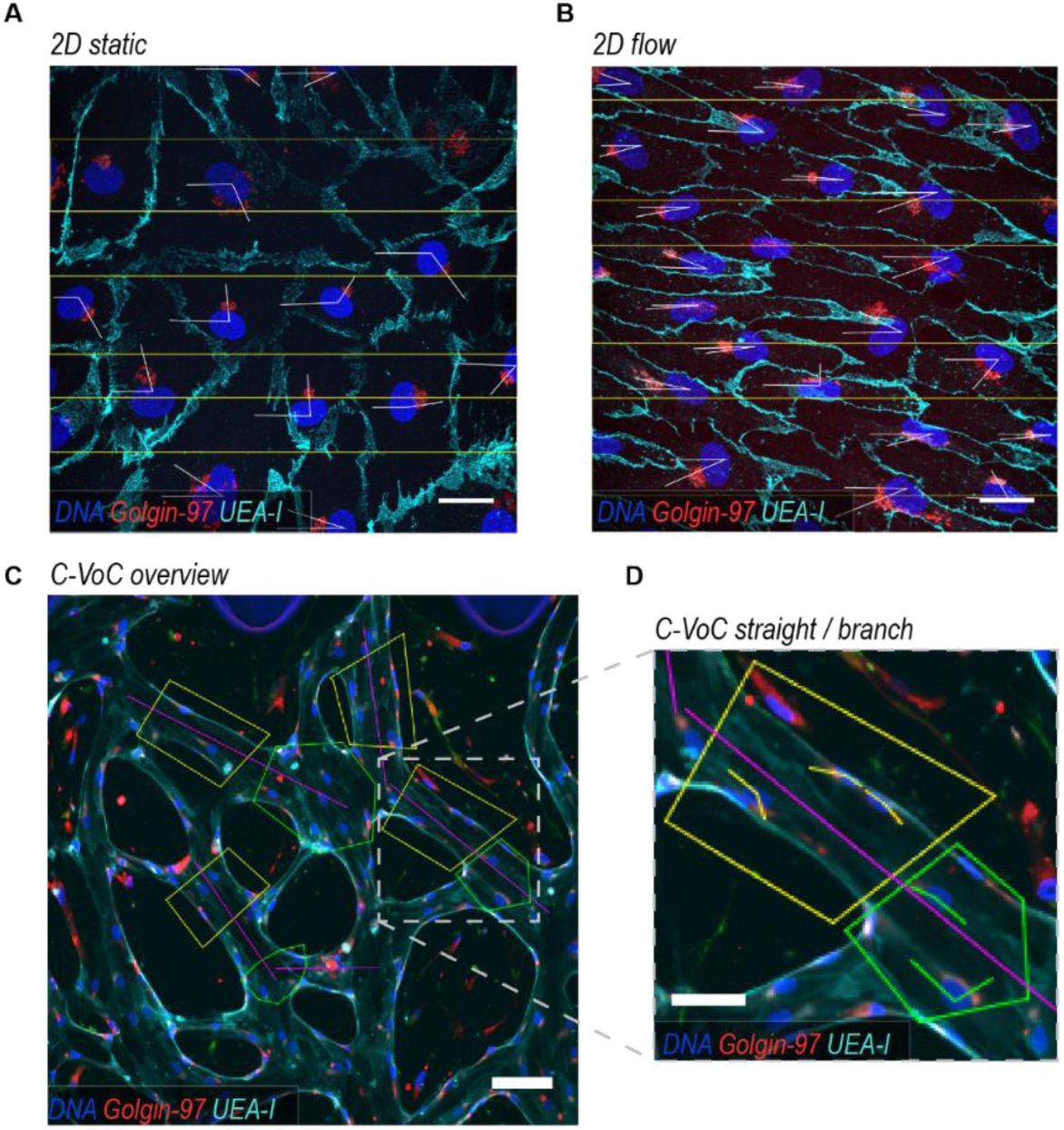
Golgi-to-nuclei angle determination gridlines and microvascular branch labeling (A) Representative confocal images of static hiPSC-ECs in 2D for Golgi-to-nuclei orientation analysis including parallel lines (yellow) as reference for golgi-to-nuclei angle determination (white angle indications) (gray: DAPI; red: Golgin – 97, cyan: VE-Cadherin). Scale bar: 25 µm. (B) Representative confocal images of hiPSC-ECs after 72h of flow in 2D for Golgi-to-nuclei orientation analysis including parallel lines (yellow) as reference for golgi-to-nuclei angle determination (white angle indications) (gray: DAPI, red: Golgin – 97, cyan: VE-Cadherin). Scale bar: 25 µm. (C) Representative confocal image of VoC for Golgi-to-nuclei orientation analysis including straight (yellow) and branched (green) labelled areas and reference line (magenta) parallel to vessel wall. (gray: DAPI, red: Golgin – 97, cyan: UEA-I). Scale bar: 100 µm. (D) Close up image (gray dashed indicated area indicated in (C)) of representative straight (yellow) and branched (green) labelled areas in VoC. Magenta line indicates reference line parallel to vessel wall. Golgi-to-nuclei angle determination parallel to reference line in the straight labelled area indicated in yellow angle, in branched areas indicated with a green angle. (gray: DAPI, red: Golgin – 97, cyan: UEA-I). Scale bars: 50 µm.

**Figure S2:**
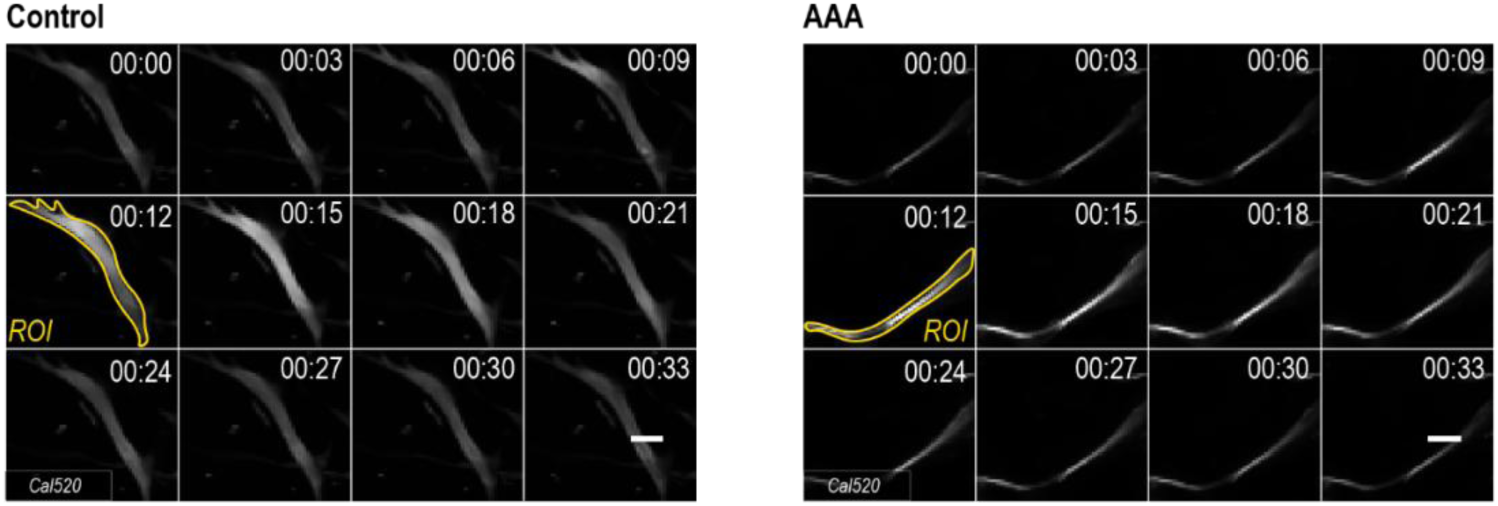
Ca2^+^ transient imaging and ROI determination Representative image for ROI detection of C-VSMC (left panel) and AAA-VoC (right panel) in VoC 5 minutes after ET-1 stimulation over the time course of 33 seconds. Representative ROI in yellow outline. (Time stamp: MM:SS, gray: Cal520). Scale bars: 25 µm.

**Figure S3:**
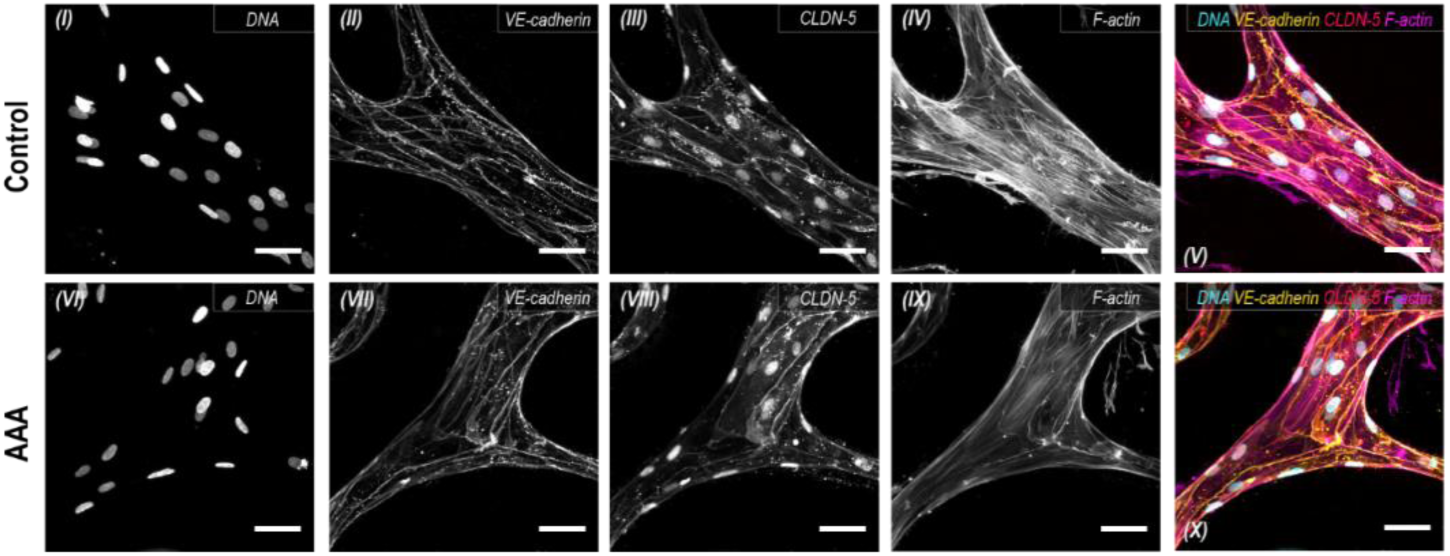
Tight and Adherens junction expression of hiPSC-ECs in Vessel-on-Chip Representative confocal images of microvascular networks, upper panel in C-VoC, lower panel in AAA-VoC. (I + VI) cell nuclei, (II + VII) VE-Cadherin, (III + VIII) CLDN-5 and (IV + IX) F-actin. (V) Composite images for C-VoC and (X) AAA-VoC (blue: DAPI, yellow: VE-Cadherin, red: CLDN-5, magenta: F-actin). Scale bars: 50 µm.

**Figure S4:**
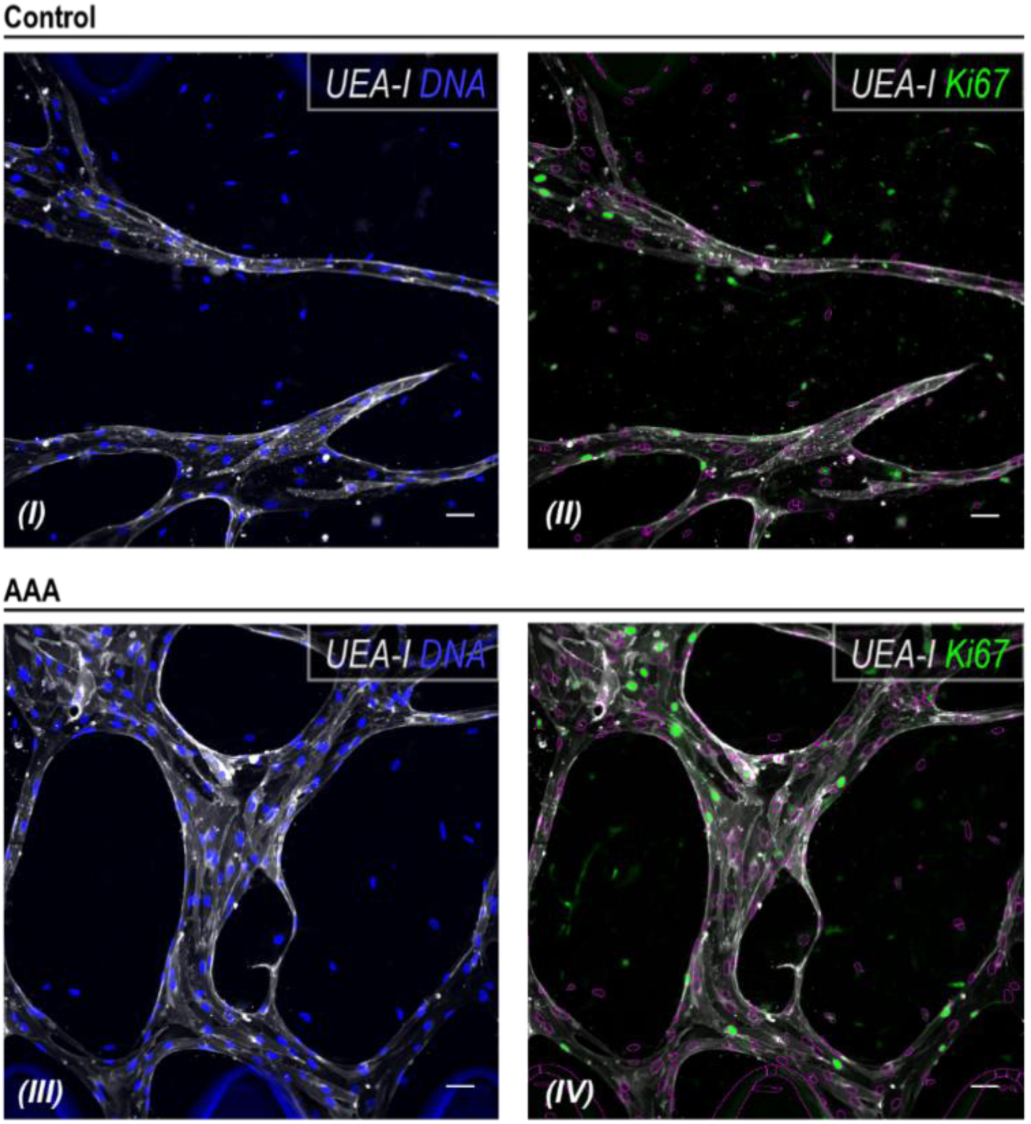
Ki67 analysis on VoC platform (I–IV) Representative confocal images of VoC for Ki67 analysis. (I) C-VoC showing nuclei for ROI determination (blue: DAPI, gray: UEA-I). (II) C-VoC with ROI boundaries (purple lines) defined by thresholding the DNA stain in (I) (green: Ki67, gray: UEA-I). (III) AAA-VoC showing nuclei for ROI determination (blue: DAPI, gray: UEA-I). (IV) AAA-VoC with ROI boundaries (purple lines) defined by thresholding the DNA stain in (III) (green: Ki67, gray: UEA-I). Scale bars: 50 µm.

**Figure S5:**
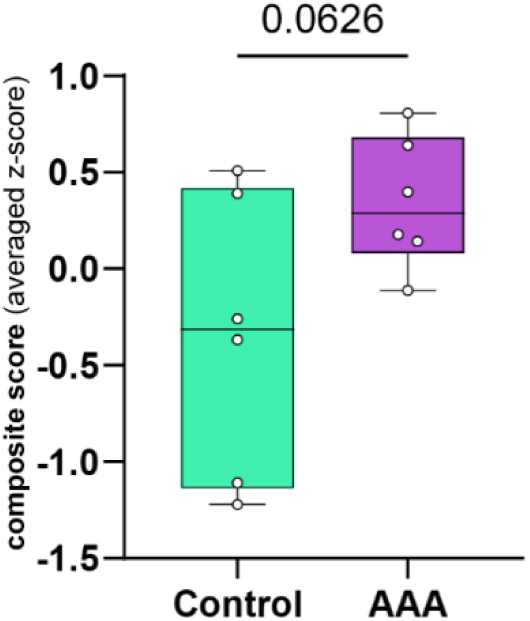
Z-scored composite score of cytokine expression Composite score of cytokine expression levels after log-transformation and z-score normalization across all samples within each group. Composite score values are averaged per sample. Data are shown as box plots: median (line), interquartile range (box), and minimum to maximum values. Each point represents an individual sample (n = 3 independent VSMC lines, 2 technical replicates each).

**Figure S6:**
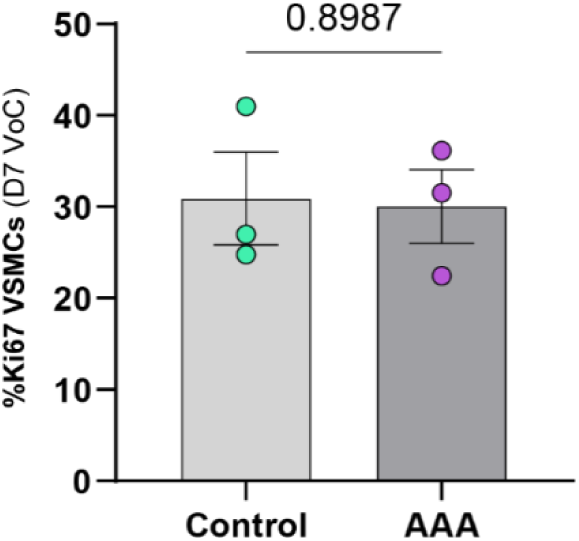
VSMC fraction of Ki67% nuclei Quantification of fraction of Ki67 positive VSMC nuclei in VoC culture at day 7 in %. Data are shown as mean ± SEM. Data points represent 3 independent experiments on 3 (control) and 3 (AAA) independent VSMC lines; 3–6 VoC channels per experiment were imaged and data averaged to yield one value per cell line. Data passed Shapiro– Wilk normality test, groups compared using unpaired two-tailed t-test. *p< 0.05.

**Figure S7:**
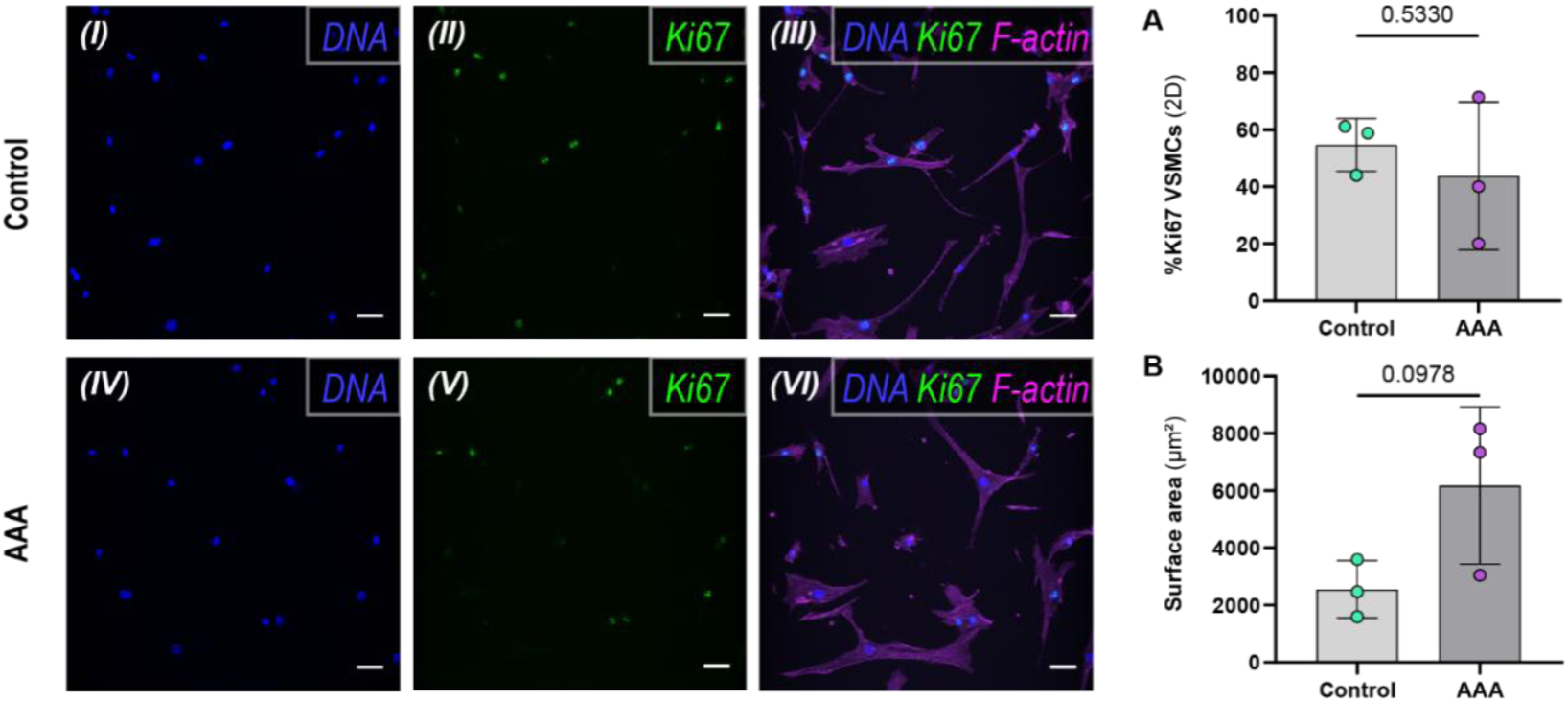
VSMC proliferation and surface area in 2D (I–IV) Representative confocal images C-VSMCs (I-III) and AAA-VSMCs (IV-VI) in 2D culture. (I + VI) cell nuclei, (II + VII) Ki67, (III + VIII) Composite (blue: DNA, green: Ki67. Magenta: F-actin). (A) Quantification of Ki67% positive cell nuclei for C-VSMCs and AAA-VSMCs. (B) Quantification of surface area in µm² for C-VSMCs and AAA-VSMCs in 2D, ROI selection based on F-actin staining on 2D maximum intensity projection images. Data are shown as mean ± SD. Data points represent 3 independent experiments on 3 (control) and 3 (AAA) independent VSMC lines, 1 image per cell line. Data passed Shapiro–Wilk normality test, groups compared using unpaired two-tailed t-test, significance level p< 0.05. Scale bars: 50 µm.

**Video S1.**
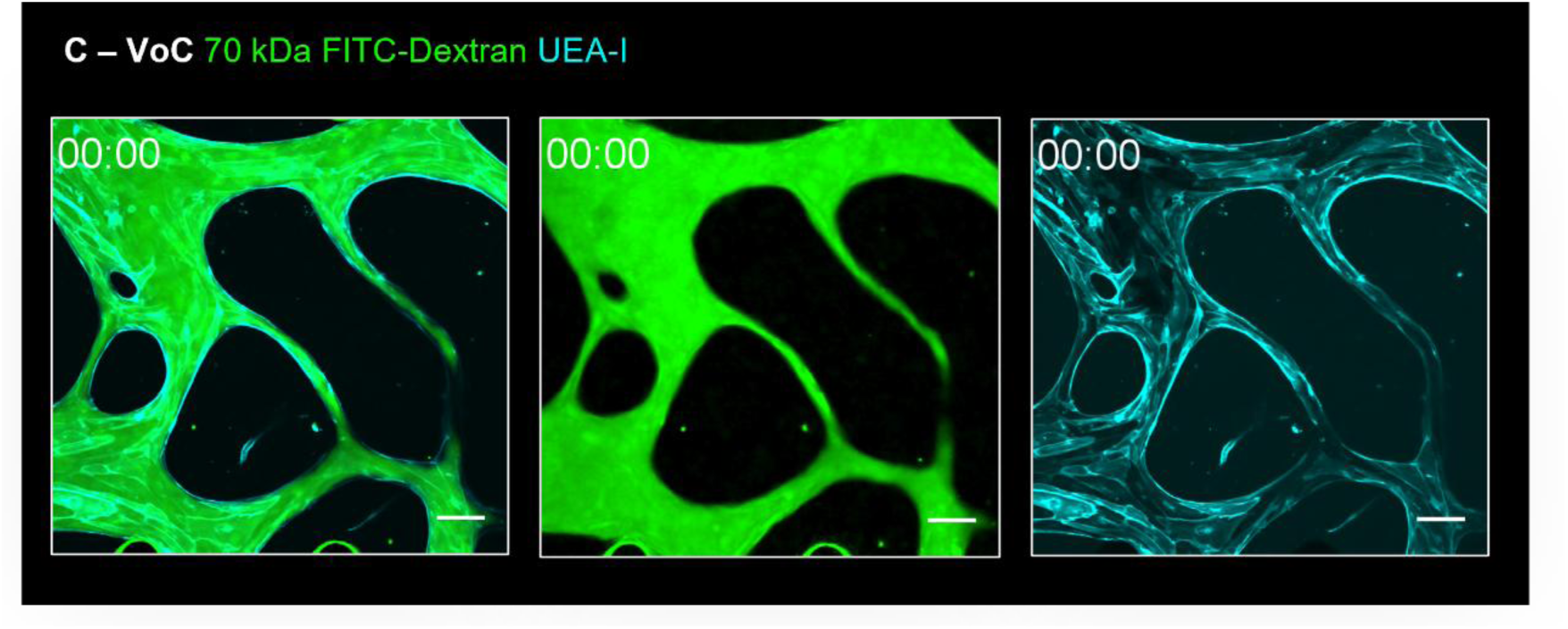
: Endothelial barrier in C-VoCs Representative video of C-VoC microvasculature (cyan: UEA-I) perfused with 70 kDa FITC-Dextran (green). Representative ROI indicated in yellow. Left: Composite (cyan: UEA-I, green: 70 kDa FITC-Dextran). Middle: 70 kDa FITC-Dextran (green). Right: Microvascular network (cyan: UEA-I). Timestamp: MM:SS. Scale bars: 100 µm.

**Video S2.**
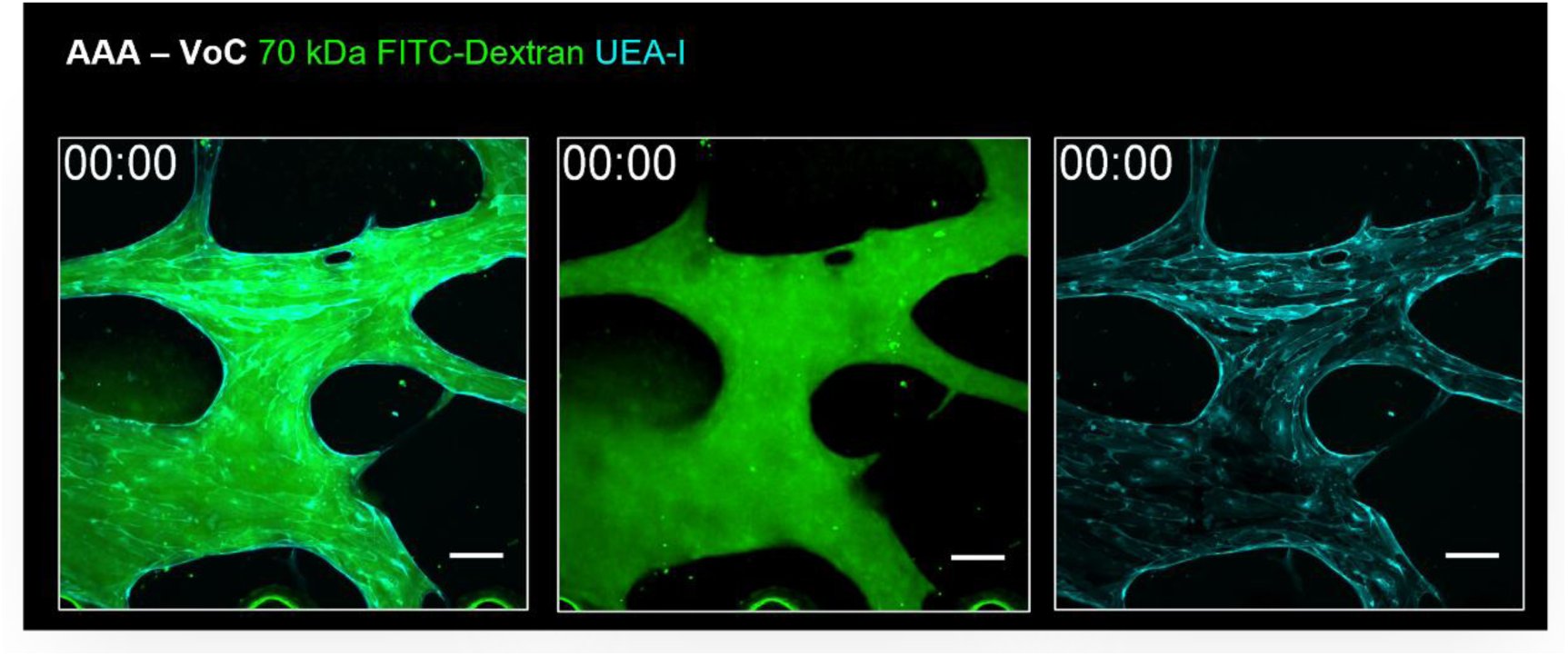
: Endothelial barrier in AAA-VoCs Representative video of C-VoC microvasculature (cyan: UEA-I) perfused with 70 kDa FITC-Dextran (green). Representative ROI indicated in yellow. Left: Composite (cyan: UEA-I, green: 70 kDa FITC-Dextran). Middle: 70 kDa FITC-Dextran (green). Right: Microvascular network (cyan: UEA-I). Timestamp: MM:SS. Scale bars: 100 µm.

